# ROMO1 loss in cholinergic neurons induces mitochondrial ultrastructural damage and ALS-like neuromuscular degeneration

**DOI:** 10.1101/2025.05.24.655877

**Authors:** Wei Nie, Zhiwen Jing, Linling Cheng, Lin-Lin Li, Fengli Xu, Limei Lin, Xiaohui Dong, Hua Han, Heping Cheng, Xianhua Wang

**Author notes:** Correspondence (Xianhua Wang); (Heping Cheng).

## Abstract

Mitochondrial dysfunction is strongly associated with the pathogenesis of amyotrophic lateral sclerosis (ALS), a neurodegenerative disorder characterized by progressive motor neuron degeneration. However, it remains obscure whether mitochondrial abnormities are sufficient to drive ALS development. Here, we show that selective depletion of reactive oxygen species modulator 1 (ROMO1), an inner mitochondrial membrane-delimited protein, in cholinergic neurons leads to adult-onset, progressive locomotor deficits in mice that closely resemble ALS pathology. ROMO1 ablation in cholinergic neurons induces ALS-like neuromuscular degeneration, as evidenced by age-dependent motor neuron loss, axon degeneration, disrupted cholinergic transmission, neuromuscular junction denervation, and the resultant muscle atrophy. Notably, ROMO1 loss induces early and progressive mitochondrial cristae deformation in motor neurons, preceding the onset of ALS-like syndromes. Our findings support that mitochondrial impairment in vulnerable motor neurons is a sufficient contributor to ALS etiology, positioning mitochondria as a potential therapeutic target.

## Introduction

Amyotrophic lateral sclerosis (ALS) is a devastating neurodegenerative disease characterized by the degeneration of both upper and lower motor neurons, leading to progressive muscle atrophy and, ultimately, paralysis ^1–3^. Approximately 10% of ALS cases are familial (fALS), predominantly autosomal dominant, while the remaining 90% are sporadic, with no clear genetic basis and variable clinical presentations ^4,5^. Given that the mutations associated with fALS are present in all tissues of the patients, it remains unclear whether ALS symptoms arise from mutation-induced dysfunction in motor neurons or in other tissues.

Motor neurons are highly polarized cells with long axons that can extend up to 1 meter. They are high-energy-demand and their ATP is largely produced through mitochondrial oxidative phosphorylation ^6,7^. Mitochondria, the double-membrane organelles critical for cellular energy production, possess a highly specialized inner membrane that invaginates into the matrix to form tubular or lamellar cristae structures, maximizing the surface area to accommodate plenty of functional proteins essential for energy metabolism ^8,9^. Morphological abnormalities in mitochondria, such as swelling or vacuolation, are commonly observed in motor neurons of both ALS patients and mouse models ^10–14^. For instance, overexpression of ALS-associated mutations like SOD1 G93A and TDP-43 A315T disrupts mitochondrial morphology, causing elongation and swelling ^15–17^ or aggregation and fragmentation ^18^, respectively. Additionally, mutation of *CHCHD10*, which encodes a mitochondrial protein critical for maintaining mitochondrial integrity and cristae junctions ^19,20^, has been identified in ALS patients ^21^, providing evidence that disrupted mitochondrial structure might contribute to ALS pathology. However, the role of *CHCHD10* mutation in ALS development remains controversial, as mice harboring this mutation also present symptoms not exclusive to motor neuron impairments, including broader skeletal and cardiac muscle defects ^22–24^. In addition to structural abnormalities, other mitochondrial dysfunctions such as aberrant mitochondrial DNA, impaired mitochondrial respiratory chain activity, and oxidative stress have been identified in the muscle and spinal cord of sporadic ALS patients ^25–31^. All these findings underscore a close interlink between mitochondrial damage and ALS progression. Yet, it remains unclear whether mitochondrial dysfunction is a consequence or a driver of ALS.

Reactive oxygen species (ROS) modulator 1 (ROMO1) is a small protein delimited in the inner mitochondrial membrane and has been implicated in various biological processes, including modulating ROS production ^32,33^ and regulating mitochondrial fusion ^34^. However, its physiological and pathological roles are poorly understood. Here, we show that selective depletion of *Romo1* in cholinergic neurons leads to ALS-like neuromuscular degeneration in mice. Notably, this degeneration is preceded by progressive mitochondrial cristae deformation in motor neurons. These findings reveal that ROMO1 is crucial for maintaining mitochondrial ultrastructure in motor neurons and that mitochondrial deformation in these neurons is sufficient to drive ALS-like pathology.

## Results

### Cholinergic neuron-selective deletion of *Romo1* induces age-dependent ALS-like locomotor deficits in mice

To investigate the physiological role of ROMO1 in motor neurons, we generated cholinergic neuron-specific *Romo1* knockout (*Romo1*-cKO) mouse model by crossing mice harboring floxed *Romo1* (*Romo1^fl/fl^*) with choline acetyltransferase (*ChAT*)*-Cre*^+/−^ mice (Figure S1A). The *Romo1*-cKO mice were viable and fertile, with offspring following expected Mendelian ratios (Figure S1B). Interestingly, *Romo1*-cKO mice developed ALS-like locomotor deficits, exhibiting overt hind limb weakness at 4 months of age (Figure 1A). Their symptoms progressively worsened to hindlimb paralysis by 5 months and extended to forelimb paralysis by 12 months of age (Figure 1A, Movie S1). In contrast, the control mice displayed normal motor activities throughout the study period (Figure 1A, Movie S1). A gradual loss of body weight was observed in both male and female *Romo1*-cKO mice, contrasting with the continuous weight gain in control mice during the measurement period (Figure 1B). Specifically, the male and female *Romo1*-cKO mice began losing weight at around 10-weeks and 14-weeks old, respectively (Figure 1B). Comparable tibia length was found between control and *Romo1*-cKO mice of 12-month old (Figure S1C), suggesting that the weight loss in *Romo1*-cKO mice is due to muscle atrophy rather than growth defects. To characterize progressive muscle weakness, we measured grip strength in control and *Romo1*-cKO mice at different ages. Before the age of 8-week, both control and *Romo1*-cKO mice showed similar increases in grip strength with age (Figure 1C), correlating to the weight gain observed during this period (Figure 1B). However, a striking and progressive muscle weakness was evidenced afterwards in both male and female *Romo1*-cKO mice, while the control mice maintained stable muscle performance during the age of 8-20 weeks (Figure 1C). Meanwhile, rotarod performance demonstrated a steady decline in *Romo1*-cKO mice starting at 8 weeks of age (Figure 1D), further confirming the age-dependent progression of motor dysfunction induced by ROMO1 loss.

**Figure 1.**
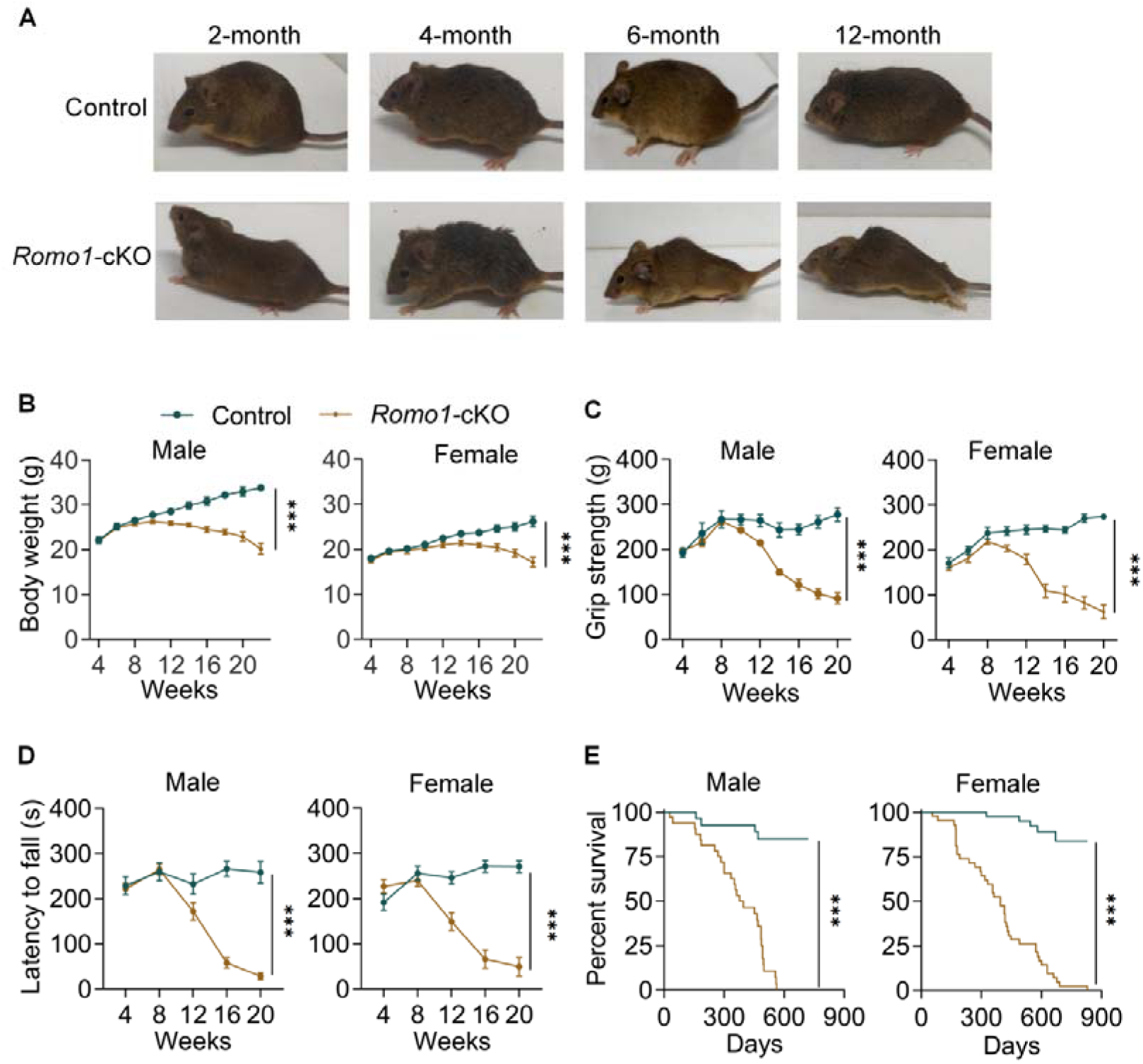
*Romo1* deletion in cholinergic neurons causes adult-onset and progressive locomotor deficits as well as shortened lifespan. (A) Photos showing that *Romo1*-cKO mice displayed progressive loss of motor capabilities and the ultimate paralysis, while the control mice exhibited normal behaviors during all the observation time. (B) Changes of body weight at different ages of control and *Romo1*-cKO mice. Data were mean ± SEM. For male mice, n=13 for control and n=19 for *Romo1*-cKO; for female mice, n=14 for control and n=19 for *Romo1*-cKO. (C) Progressive decline of grip strength after peaking at 8 weeks in *Romo1*-cKO mice. Data were mean ± SEM. For male mice, n=7 for control and n=11 for *Romo1*-cKO; for female mice, n=8 for control and n=11 for *Romo1*-cKO. (D) Rotarod tests showing a progressive decline starting from 8 weeks old in *Romo1*-cKO mice. Data were mean ± SEM. For male mice, n=7 for control and n=11 for *Romo1*-cKO; for female mice, n=8 for control and n=11 for *Romo1*-cKO. (E) Kaplan-Meier survival curves showing shorted lifespan of *Romo1*-cKO mice. For male mice, n=27 for control and n=32 for *Romo1*-cKO; for female mice, n=40 for control and n=42 for *Romo1*-cKO. For B-D, two-way ANOVA with Sidak’s multiple comparison test was used; for E, Mantel-Cox log-rank analysis was used. ***p < 0.001. The color representation of mouse genotype in Figure B–E is shown in B.

Moreover, *Romo1*-cKO mice exhibited a significant shortened lifespan, with a median lifespan of 386 days and 396 days for male and female mice, respectively (Figure 1E). The survival rates of control mice were as high as 84% and 83% by the end of their observation period of 710 days for male mice and 830 days for female mice (Figure 1E). Notably, mating and nursing behavior shortened the lifespan of female *Romo1*-cKO mice with median lifespan being 157 days (Figure S1D).

Collectively, these results reveal that loss of ROMO1 in cholinergic neurons induces an ALS-like phenotype in mice, characterized by progressive muscle weakness, locomotor deficits, and shortened lifespan, highlighting an essential role of ROMO1 in these neurons.

### ROMO1 ablation induces structural and functional impairment of motor neurons

Given that ROMO1 is selectively depleted in cholinergic neurons, we hypothesized that the observed ALS-like locomotor deficits in *Romo1*-cKO mice might be due to motor neuron damage. We then first quantified motor neurons in the ventral horn of the level 5 lumbar spinal cord by staining for ChAT-positive neurons. At 6 weeks of age, *Romo1*-cKO and control mice showed comparable motor neuron numbers (Figure 2B), suggesting that ROMO1 ablation does not affect motor neuron differentiation. However, at 8 weeks of age, when *Romo1*-cKO mice still exhibited normal locomotor activities (Figure 1C, 1D), a significant reduction in motor neurons was evident in *Romo1*-cKO mice compared to controls (Figure 2B). This loss of motor neurons was progressively worsened with age, and by 12 months, approximately half of the motor neurons had been lost (Figure 2A, 2B), coinciding with the onset of severe paralysis (Figure 1A). This gradual decline of motor neurons was further confirmed with Nissl staining (Figure S2A, S2B). Additionally, gliosis, a hallmark of neurodegeneration ^35^, was significantly elevated in 8-week old *Romo1*-cKO mice (Figure S2C, S2D), indicating an ongoing neuroinflammatory response.

**Figure 2.**
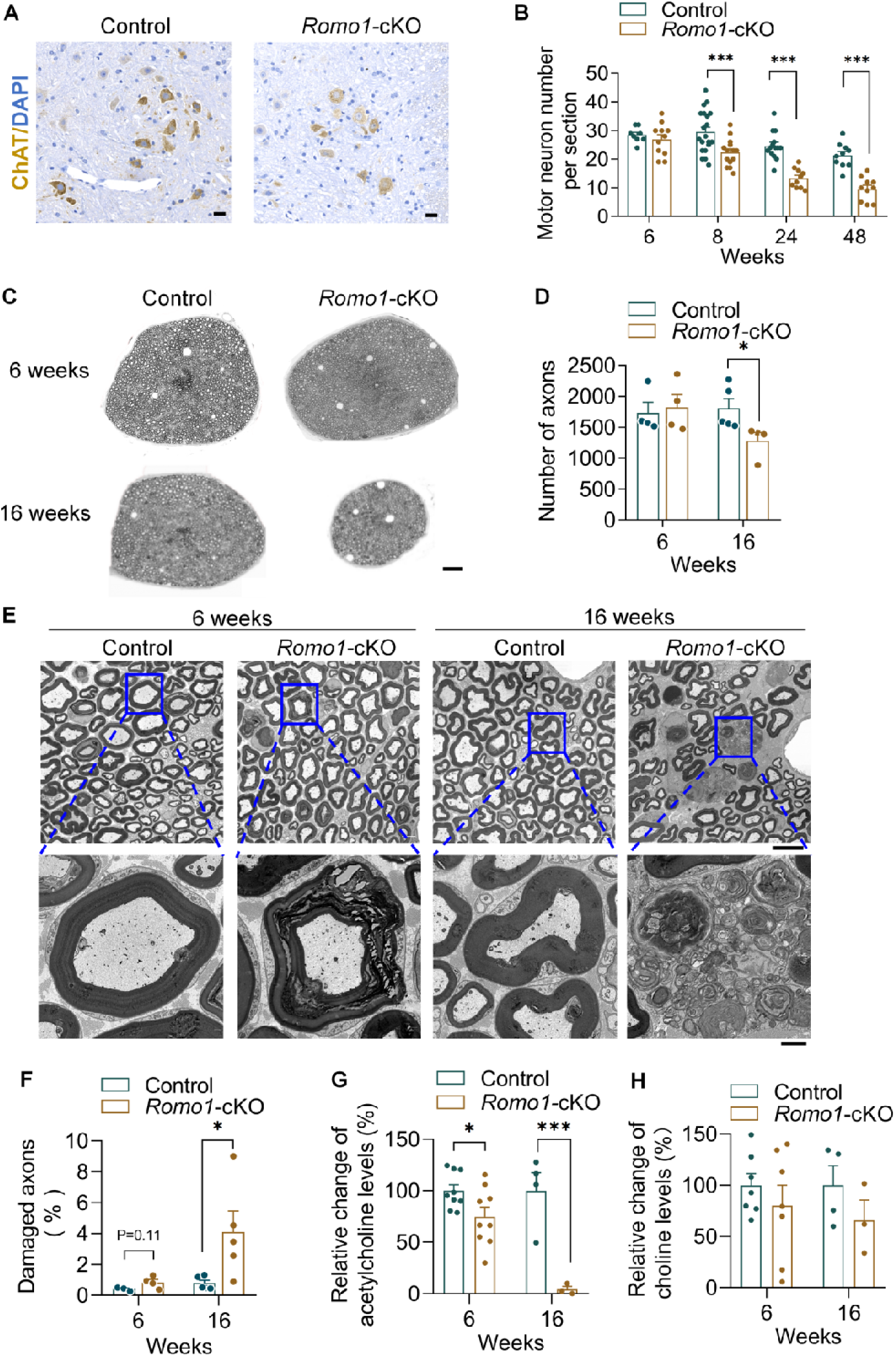
Progressive spinal motor neuron degeneration and sciatic nerve axon damage in *Romo1*-cKO mice. (A) Representative anti-ChAT immunohistochemistry for motor neurons in the ventral horn of level 5 lumbar spinal cord isolated from *Romo1*-cKO and control mice at 48 weeks. Scale bars: 20 µm. (B) Quantification of motor neuron numbers in the level 5 lumbar spinal cord from different ages of control and *Romo1*-cKO mice. Data were mean ± SEM. n=8-20 slices from 3-6 mice for each group. (C) Representative electron micrographs of cross sections of sciatic nerve from control and *Romo1*-cKO mice at 6- and 16-week old. Scale bar: 50 µm. (D) Quantification of the axon numbers in (C). Data were mean ± SEM. n=4-5 mice for each group. (E) Representative electron micrographs showing damaged axons in *Romo1*-cKO mice. Scale bars: 10 μm (top panels), 1.5 μm (bottom panels). (F) Quantification of damaged axon proportions in control and *Romo1*-cKO mice. Data were mean ± SEM. n=3-5 mice for each group. (G) Decreased acetylcholine levels in tibialis anterior muscle of *Romo1*-cKO mice. Data were mean ± SEM. n=3-9 mice for each group. (H) Comparable choline levels in tibialis anterior muscle of control and *Romo1*-cKO mice. Data were mean ± SEM. n=3-7 mice for each group. For B, D, F-H, unpaired t test was used. *p < 0.05, ***p < 0.001.

In a parallel experiment, we assessed the axonal integrity of the sciatic nerve in 6-week and 16-week old mice. At 6 weeks, there were no significant differences in axon number between *Romo1*-cKO and control mice, but by 16 weeks, a substantial reduction in axon numbers was observed in *Romo1*-cKO mice (Figure 2C, 2D). Notably, axon damage shown by myelin injury, was evident as early as 6 weeks in *Romo1*-cKO mice, and such axon damage was exacerbated with age as a larger number of abnormal axons and more severe injuries were found in older mice (Figure 2E, 2F). To further evaluate motor neuron function, we measured acetylcholine levels in the tibialis anterior muscle. As early as 6 weeks, acetylcholine levels were significantly reduced in *Romo1*-cKO mice compared to controls (Figure 2G), with the levels almost undetectable by 16 weeks (Figure 2G). Choline levels were comparable between *Romo1*-cKO and control mice at both time points (Figure 2H). These results underscore an important role of ROMO1 in maintaining structural and functional integrity of motor neurons.

### ROMO1 ablation in cholinergic neurons leads to progressive denervation of the neuromuscular junction, resulting in muscle atrophy

A hallmark of ALS pathology is denervation at the synapse between the nerve and muscle, known as the neuromuscular junction (NMJ) ^36,37^. To examine whether ROMO1 ablation affects NMJ integrity, we first assessed the structural integrity of the NMJ in *Romo1*-cKO and control mice at different ages. At 6 weeks, prior to noticeable muscle weakness, both control and *Romo1*-cKO mice displayed intact NMJs, with clustered acetylcholine receptors forming characteristic postsynaptic endplates which are consistently colocalized with presynaptic terminals (Figure 3A, 3B). However, by 8 weeks of age, when *Romo1*-cKO mice began showing significant motor neuron degeneration (Figure 2), NMJs in *Romo1*-cKO mice were severely disorganized, with only about 40% of the endplates still innervated (Figure 3A, 3B). By 24 weeks when *Romo1*-cKO mice developed severe locomotor deficits and motor neuron degeneration (Figure 1, 2), the NMJ denervation was exacerbated further, leaving fewer than 4% of the endplates innervated, in stark contrast to 100% innervation in control mice (Figure 3A, 3B). Concurrently, a significant reduction in acetylcholine receptor density at the postsynaptic membrane was also observed in *Romo1*-cKO mice of 24-week old (Figure 3A), likely a consequence of persistent NMJ denervation.

**Figure 3.**
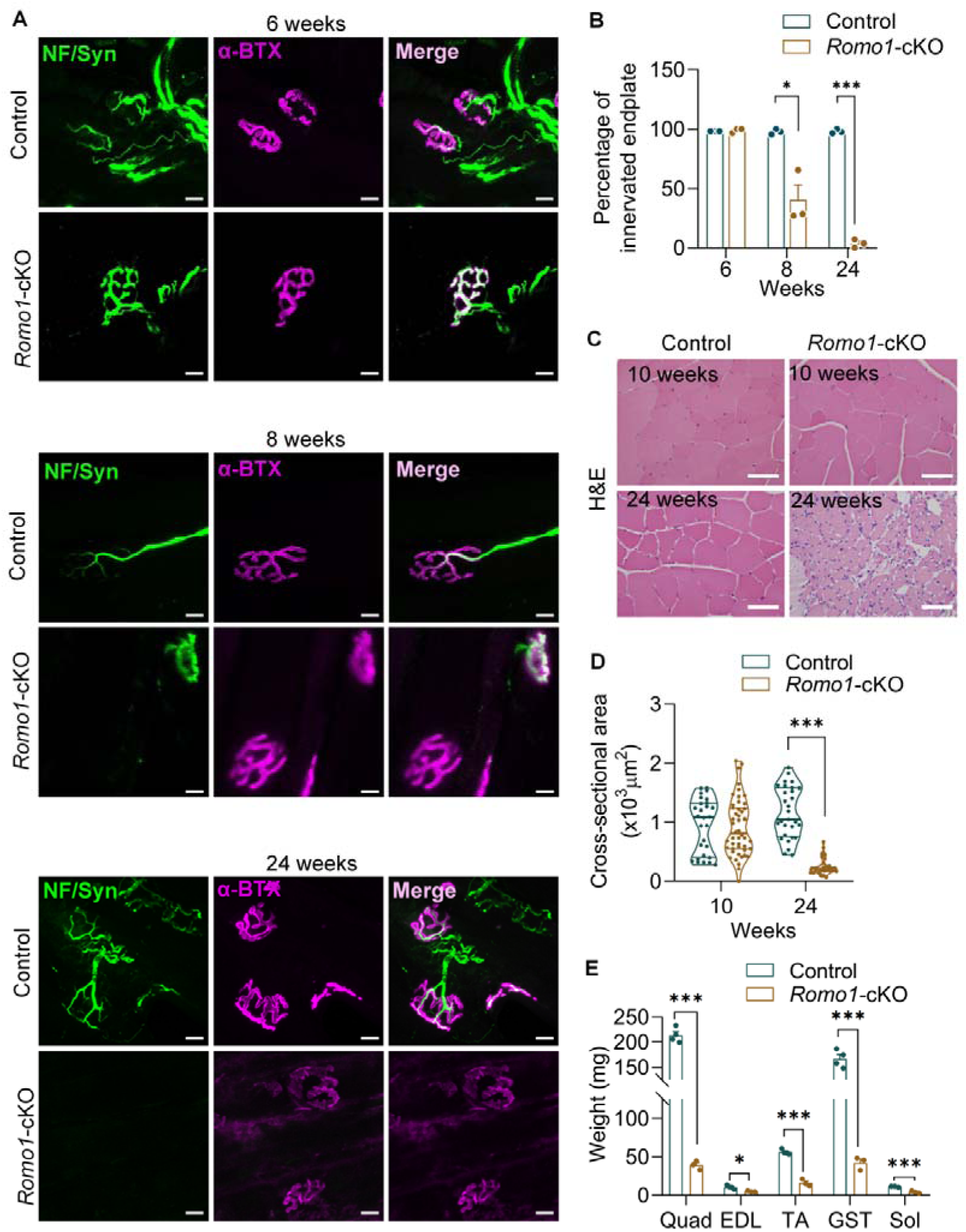
Progressive NMJ denervation and the resultant muscle atrophy in *Romo1-*cKO mice. (A) Representative confocal images of innervated and denervated endplates of the tibialis anterior muscle in *Romo1*-cKO and control mice at 6, 8 and 24 weeks old. Anti-neurofilament (NF) and anti-synaptophysin (Syn) label presynaptic motor neuron projections and α-Bungarotoxin (α-BTX) labels postsynaptic acetylcholine receptors. Scale bars: 10 μm. (B) Quantification of muscle denervation in tibialis anterior muscles. Data were mean ± SEM. n=3 mice for each group. (C) Representative hematoxylin and eosin (H&E) staining images of cross sections of tibialis anterior muscles isolated from *Romo1*-cKO and control mice. Scale bars: 40 μm. (D) Quantification of muscle cross-sectional fiber area in *Romo1*-cKO and control mice. Data were mean ± SEM. n=28-48 cells from 3 mice for each group. (E) Reduction of muscle mass in *Romo1*-cKO mice at 48 weeks old. Data were mean ± SEM. n=3-4 mice for each group. For B and E, unpaired t test was used; for D, unpaired, two-tailed Mann-Whitney U-Test was used. *p < 0.05, ***p < 0.001.

We next determined whether progressive NMJ denervation led to muscle atrophy in *Romo1*-cKO mice. Histological examination of tibialis anterior muscle showed that muscle fibers in *Romo1*-cKO mice were structurally normal at 10 weeks of age (Figure 3C, 3D), when the motor neurons were already damaged (Figure 2). However, by 24 weeks, a dramatic atrophy of muscle fibers was evident in *Romo1*-cKO mice (Figure 3C, 3D). Notably, no significant infiltration of inflammatory cells was detected in the muscle tissue (Figure 3C), suggesting that the muscle atrophy observed was neurogenic, rather than resulting from myogenic necrosis. At 12 months, when *Romo1*-cKO mice had developed complete paralysis (Figure 1), they exhibited consistent and severe muscle atrophy across all muscle types measured, including the quadriceps, extensor digitorum longus, tibialis anterior, gastrocnemius and soleus (Figure 3E). The post onset of muscle atrophy in *Romo1*-cKO mice support that it is a consequence of motor neuron degeneration and NMJ denervation.

### ROMO1 loss induces early and progressive abnormalities of mitochondrial ultrastructure in motor neurons

The above results revealed that genetic ablation of *Romo1* in cholinergic neuron is sufficient to induce ALS-like pathologies in mice. Given that ROMO1 is an inner mitochondrial membrane-delimited protein, we hypothesized that the observed motor neuron degeneration in *Romo1*-cKO mice could be driven by mitochondrial dysregulation. To test this, we examined mitochondrial ultrastructure in the soma of spinal cord motor neurons from control and *Romo1*-cKO mice at 6-week and 16-week old. Mitochondria were similarly dispersed around the soma in both control and *Romo1*-cKO motor neurons from 6-week old mice (Figure 4A). However, by 16 weeks, mitochondrial distribution in *Romo1*-cKO motor neurons became severely disrupted, with mitochondria aggregating at the cell periphery in some motor neurons or being lost in the other part of cells (Figure 4A). Upon closer examination, we observed severe mitochondrial cristae deformation in *Romo1*-cKO mice as early as 6 weeks (Figure 4B), when the motor neuron soma displayed no overt histopathological lesions (Figure 2B), highlighting early onset of mitochondrial ultrastructural dysregulation induced by ROMO1 deficiency. The mitochondrial abnormalities in motor neurons were exacerbated in 16-week old mice as evidenced by severely worsened cristae deformation (Figure 4B). To quantitatively assess the degree of mitochondrial abnormalities, we classified mitochondria into four distinct types based on their cristae structure: type I with normal lamellar cristae, type II with swollen cristae, type III with sparse and swollen cristae, and type IV with concentric cristae (Figure 4B). Comparing to the normal type I mitochondria, type II and III were enlarged, suggesting mitochondrial swelling, whereas type IV exhibited a shrunken area indicative of fragmentation (Figure 4C). At 6 weeks of age, most of the mitochondria in *Romo1*-cKO cells were type II (∼89%) and type III (∼9%), while at 16 weeks, there was a significant shift toward type III (∼40%) and type IV (∼58%) (Figure 4D). In contrast, ∼99% of mitochondria in control motor neuron soma were type I at both ages (Figure 4D). The mitochondria in *Romo1*-cKO cells exhibited greater circularity at 6 weeks (Figure 4E), substantiating increased swelling in type II and III mitochondria. These results indicate that ROMO1 is required for maintaining normal mitochondrial ultrastructure in motor neurons.

**Figure 4.**
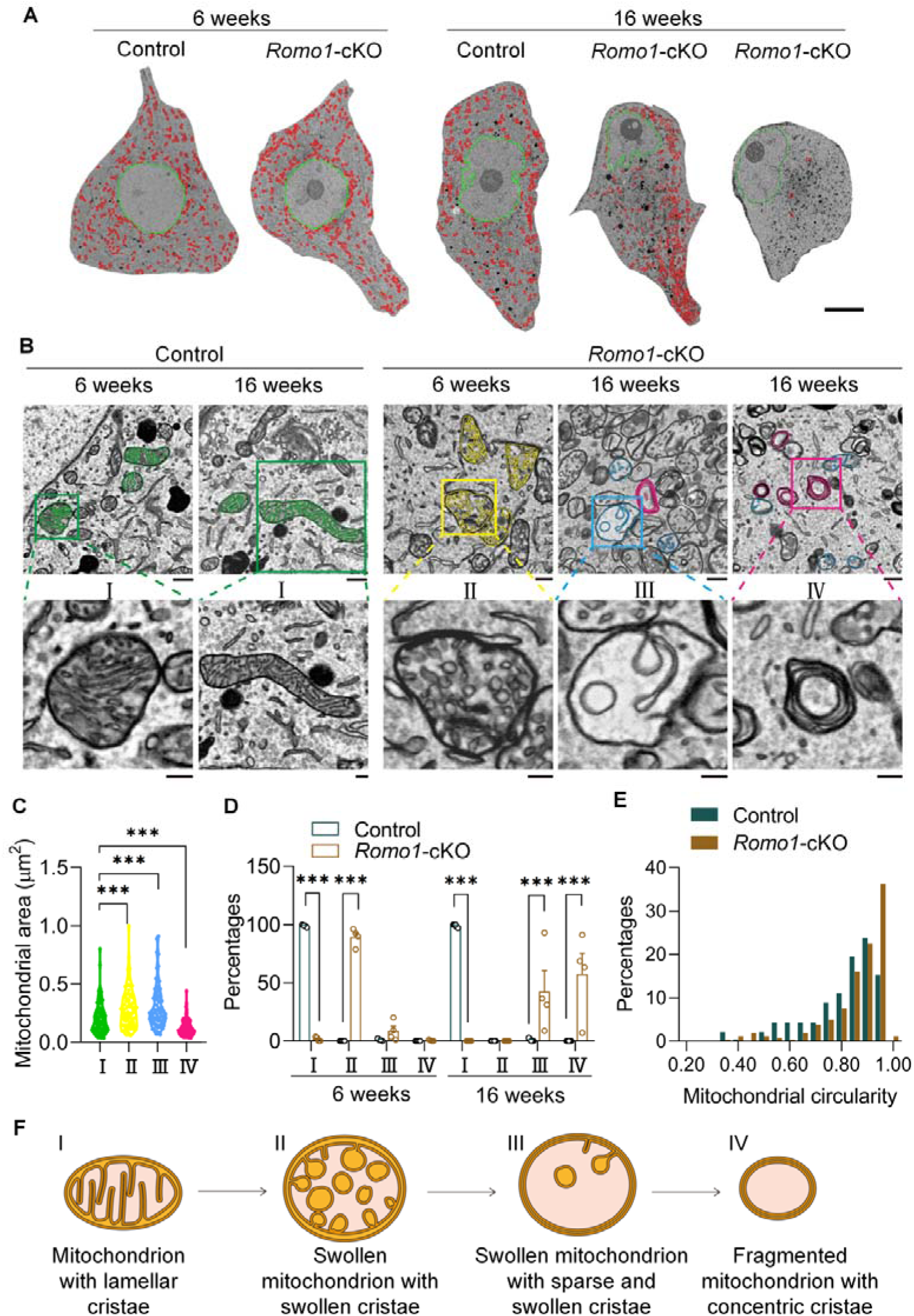
ROMO1 loss induces mitochondrial ultrastructural aberrance with cristae deformation in motor neurons. (A) Representative electron micrographs of motor neuron soma in the level 5 lumbar spinal cord showing abnormal mitochondrial distribution or loss in 16-week old *Romo1*-cKO mice. The nuclei and mitochondria are marked in green and red, respectively. Scale bar: 10 μm. (B) Representative electron micrographs showing deformation of mitochondrial cristae in *Romo1*-cKO motor neurons. In specific, mitochondria were divided into four types according to their cristae morphology: mitochondria with typical lamellar cristae (I), mitochondria with swollen cristae (II), mitochondria with sparse and swollen cristae (III), and mitochondria with concentric cristae (IV). Scale bars: 500 nm (top panels), 200 nm (bottom panels). (C) Area analysis of the four types of mitochondria. n=80 mitochondria from 3 mice for each group. (D) Percentages of the four different mitochondrial types in *Romo1*-cKO and control motor neurons. Data were mean ± SEM. n=3-5 neurons for each group. (E) Percentage distribution of mitochondrial circularity in motor neurons of 6-week old *Romo1*-cKO and control mice. n=236-242 mitochondria from 3 mice for each group. (F) Diagram illustrating progressive deformation of mitochondrial cristae in ROMO1-deficient motor neurons. For C, unpaired, two-tailed Mann-Whitney U-Test was used; for D, two-way ANOVA with Sidak’s multiple comparison test was used. ***p < 0.001.

Notably, mitochondria in adjacent non-motor neurons within *Romo1*-cKO spinal cord sections retained their normal morphology, showing typical lamellar cristae (Figure S3A), indicating a selective deleterious impact on mitochondrial ultrastructure in this mouse model and suggesting that motor neuron degeneration caused by mitochondrial dysregulation is sufficient to cause ALS-like pathology. Additionally, the endoplasmic reticulum (ER) of *Romo1*-cKO motor neurons exhibited mild swelling at the age of 6 weeks, which progressed into more severe fragmentation and loss of ER architecture by 16 weeks (Figure S3B). This ER aberrance is likely a secondary consequence of mitochondrial disruption.

To investigate the functional impact of ROMO1 loss, we utilized the motoneuron-like NSC34 cell line ^38^ for further analysis. In *Romo1* knockdown NSC34 cells, we observed a consistent reduction in intracellular acetylcholine levels without changes in choline levels (Figure S4A-S4C). The mitochondria in *Romo1* knockdown cells appeared more rounded (Figure S4D, S4E), smaller (Figure S4F), and contained fewer cristae (Figure S4G), mirroring the mitochondrial alterations observed in *Romo1*-cKO motor neurons (Figure 4), though to a lesser extent. Moreover, *Romo1* knockdown led to a decrease in cellular ATP contents (Figure S4H) and an increase in ROS levels (Figure S4I), supporting that ROMO1 loss impairs mitochondrial functions.

Collectively, these results indicate that ROMO1 ablation in motor neurons leads to early, progressive damages to mitochondrial ultrastructure. Initially, mitochondria swell with swollen cristae, followed by reduction of cristae, and eventually fragmentation with concentrically deformed cristae (Figure 4F). The early onset of mitochondrial ultrastructural aberrance, preceding motor neuron degeneration, points to a casual role of mitochondrial damage in the development of ALS-like pathologies in *Romo1*-cKO mice.

## Discussion

In this study, we demonstrate that selective ablation of ROMO1 in cholinergic neurons leads to an ALS-like syndrome, characterized by adult onset and progressive neuromuscular degeneration. Our results reveal that ROMO1 loss induces early and age-dependent mitochondrial ultrastructural disruptions in motor neurons, which precedes the onset of motor neuron degeneration. These findings suggest that mitochondrial dysregulation is a critical driver of motor neuron damage, ultimately leading to ALS-like pathology. Our work underscores a critical role of ROMO1 in maintaining mitochondrial structural and functional integrity in motor neurons and provide strong evidence that mitochondrial injury in these vulnerable cells substantially contributes to ALS pathogenesis.

While a hallmark of ALS pathologies is motor neuron damage and consequential skeletal muscle atrophy ^39–41^, it remains obscure whether the defects in motor neurons are the primary cause of ALS. Here, we show that selective ablation of ROMO1 in cholinergic neurons induces progressive motor neuron damage that leads to ALS-like neuromuscular degeneration. Our results support that motor neuron-originated defects drive ALS pathology, though it is important to acknowledge the potential role of other cell types affected by motor neuron degeneration as secondary contributors to disease progression.

Mitochondrial dysfunction is a well-known feature of ALS ^42,43^, but only one mitochondrial protein CHCHD10 has been implicated in fALS ^21^. Whether mitochondrial dysregulation is a consequence or a driver of ALS pathologies is still an open question. In our study, we show that ablation of ROMO1, an inner mitochondrial membrane protein, leads to mitochondrial ultrastructural aberrance in motor neurons, with striking and gradual cristae deformation. Importantly, these mitochondrial structural damages precede of the onset of ALS-like symptoms in this *Romo1*-cKO mouse model, supporting the idea that ROMO1 loss-induced mitochondrial dysregulation is a cause rather than a consequence of ALS-like pathologies. Particularly, at young age of 6-week, when *Romo1*-cKO mice still display normal locomotor activities, they already show mitochondrial cristae deformation and lowered acetylcholine levels, suggesting that mitochondrial damage disrupts cholinergic transmission in motor neurons before overt motor neuron degeneration occurs. This aligns with previous findings showing reduced ChAT activity in single spinal motor neurons of ALS patients early in disease ^44^, as well as decreased ChAT level prior to motor neuron loss and NMJ detachment in ALS mouse models ^45^. Future studies are required to explore how ROMO1 loss leads to disrupted cholinergic transmission in motor neurons. Nonetheless, our findings strongly support the idea that mitochondrial dysfunction in motor neurons contributes significantly to ALS etiology.

ROMO1 have been shown to modulate cristae shape by influencing OPA1 oligomerization ^34^. In *Romo1* knockdown U2OS cells, mitochondrial cristae are reduced or absent, with a small subset of cells displaying cristae stacks ^34^. In our study, we found that ROMO1 ablation in motor neurons induces severe and progressive cristae deformation, starting with swollen cristae, followed by cristae loss and ultimately the formation of aberrant concentric structures. Along with the cristae deformation, mitochondria undergo swelling and then fragmentation in ROMO1 deficient motor neurons. Future studies await to elucidate the molecular mechanisms underlying ROMO1 loss-induced mitochondrial ultrastructural changes, particularly whether ROMO1 ablation impairs OPA1 oligomerization or affects other mitochondrial membrane organization players.

A major limitation in ALS research is the lack of proper animal models that accurately represent ALS pathologies. Most current models are based on genetic mutations identified in fALS, which fail to replicate the complex etiology of sporadic ALS ^4^. The *Romo1*-cKO mouse model constructed in this study replicates key features of ALS, including adult-onset motor neuron degeneration, progressive motor weakness, and eventually paralysis and death. This mouse model provides a valuable new tool for studying ALS pathogenesis, particularly in the context of sporadic ALS. Further research is warranted to determine the relevance of *ROMO1* mutations in sporadic ALS and to investigate how ROMO1 dysfunction contributes to the onset and progression of this neurodegenerative disease.

## Material and Methods

### Animals

All animal procedures were conducted in accordance with guidelines from the Association for Assessment and Accreditation of Laboratory Animal Care (AAALAC) and the Guide for the Care and Use of Laboratory Animals published by the US National Institutes of Health (NIH Publication eighth edition, update 2011). All procedures were approved by the Animal Care Committee of Peking University (IMM-ChengHP-1, 14). All mice were group housed with food and water available ad libitum in a 12-hour light/dark environment at 20-22L and 30-70% humidity.

### Generation of cholinergic neuron-selective *Romo1* knockout mouse model

Floxed *Romo1* mice (*Romo1^fl/fl^,* C57BL/6J) were generated by standard techniques using a targeting vector containing a neomycin resistance cassette flanked by FRT sites. To generate cholinergic neuron-specific *Romo1* knockout mouse model, *Romo1^fl^*^/*fl*^ mice were crossed with *ChAT-Cre^+/-^* transgenic mice (B6; 129S6-Chattm2(cre)Lowl/J, JAX stock 006410), in which Cre recombinase is expressed in cholinergic neurons under the control of the *ChAT* promoter without disrupting endogenous *ChAT* expression, to get *Romo1^fl/+^*-*ChAT-Cre^+/-^*mice. Then the heterozygous knockouts were intercrossed to generate homozygous *Romo1^fl/fl^* -*ChAT-Cre^+/-^* mice. *Romo1^fl/fl^ -ChAT-Cre^-/-^* mice were used as controls and *Romo1^fl/fl^ -ChAT-Cre^+/-^* mice were referred to *Romo1*-cKO mice. PCR was performed to confirm the genotypes.

### Grip strength test

Grip strength of fore and hind limbs was assessed using a grip strength meter (SA417, SansBio). The mice of different ages were lifted by their tails and allowed to grasp the steel grid attached to the apparatus with their forepaws and backpaws. Mice were then gently pulled across the steel grid until their grip was released. Peak tension was recorded for each of the three trials, and the best performance was recorded as the grip strength on the trial day. Mice were allowed to rest for at least 10 minutes between the trials.

### Accelerating rotarod test

Mice were first trained on the rotarod for three sessions 1 week prior to the initial analysis. For testing, mice were tested for the time on an accelerating rotarod (linear ramp from 4 rpm to 44 rpm in 5 minutes) at different ages. Three trials were administered in each test. The best performance among the three trials was recorded as the performance at each age point. Mice were allowed to have rest of at least 60 minutes between the trials.

### Immunostaining and Nissl staining of level 5 spinal cord

Mice were anesthetized with 1.25% tribromoethanol (20 mL/kg, i.p.) and transcardially perfused with phosphate-buffered saline (PBS) followed by 4% paraformaldehyde (PFA) in PBS. The level 5 spinal cord tissues were dissected and fixed in 4% PFA for 24 hours and processed for paraffin embedding and sectioning. Prior to antigen retrieval, transverse spinal cord sections (4 μm thick) were deparaffinized and rehydrated in 0.1 M citrate buffer (pH 6.0) or EDTA antigen retrieval solution (C1034, Solarbio). Sections were quenched by using 3% H_2_O_2_ and blocked in 10% normal goat serum at room temperature prior to incubation with primary antibodies. For immunohistochemistry of ChAT, the sections were incubated of recombinant monoclonal anti-ChAT antibody (ZRB1012, Sigma, 1:100) overnight at 4L followed by washing 5 minutes for three times, and then stained using the diaminobenzidine peroxidase substrate kit (ZLI-9018, ZSGB-BIO) according to the manufacturer’s instructions. The sections were then mounted with mounting medium and DAPI (ZLI-9557, ZSGB-BIO), and sealed for imaging. For Nissl staining, the 4-μm thick transverse spinal cord sections were stained with Nissl staining solution (DK0020, LEAGENE) according to the manufacturer’s instructions. Images of ChAT immunohistochemistry or Nissl staining were captured using a brightfield slide scanner (ZEISS Axio Scan. Z1). ChAT-positive motor neurons and Nissl staining-indicated big neurons in the ventral horn region were quantified from at least two different sections from each mouse. For immunofluorescence of GFAP, the sections were incubated with polyclonal anti-GFAP antibody (MAB3402, Millipore, 1:100) overnight at 4L followed by washing 5 minutes for three times, and then incubated with goat anti-rabbit Alexa 488 secondary antibody (A-11008, Molecular Probes, 1:1000) for 30Lminutes at room temperature. Images were captured by Zeiss LSM980 confocal microscope. The signal intensity of all the images was analyzed using ImageJ software (US National Institutes of Health).

### Hematoxylin and eosin staining

The excised muscles of the rear leg were fixed with 4% PFA, dehydrated, embedded in paraffin, and serially sectioned at the thickness of 4 μm. The muscle slices were mounted on a slide and deparaffinized, then hydrated using decreasing concentrations of ethanol and water. After differentiation and bluing, sections were incubated with hematoxylin for 6 minutes, gently rinsed in water, and stained in eosin for 3 minutes. Sections were rinsed in distilled water, dehydrated, and mounted. Images were captured using a brightfield slide scanner (ZEISS Axio Scan. Z1). The cross-sectional area of the fibers was analyzed using Image J software (US National Institutes of Health).

### NMJ innervation analysis

Tibialis anterior muscles were dissected and flash frozen by covering the muscle in OCT compound (4583, Sakura) and cooled in liquid nitrogen. 35-μm thick sections were acquired by a Leica cryostat. Sections were permeabilized with 0.5% Triton X-100 for 20 minutes followed by blocking with 10% horse serum in PBS for 1 hour at room temperature. Sections were stained with a rabbit anti-neurofilament heavy polypeptide antibody (C28E10, Cell Signaling Technology, 1:100) and a rabbit anti-synaptophysin (ab32127, Abcam, 1:400) overnight at 4°C. After three washes with PBS, sections were then stained with Alexa 488 nm-labeled goat anti-rabbit (A-11008, Thermo Fisher Scientific, 1:500) as well as α-bungarotoxin-tetramethylrhodamine (T0195, Sigma, 2 μg/mL) overnight at 4°C in the dark. After three washes with PBS, slides were mounted and sealed for imaging. Images were acquired on a Zeiss LSM980 confocal microscope. NMJs were selected randomly using the α-bungarotoxin signal.

### Electron microscopy analysis

Mice were anesthetized with 1.25% tribromoethanol (20 mL/kg, i.p.) and perfused at room temperature with 0.1 M phosphate buffer (PB), followed by a fixative solution (2% PFA (P6148, Sigma), 1.25% glutaraldehyde (G5882, Sigma) in 0.1 M PB). Dissected lumbar level 5 spinal cord segments or sciatic nerves were postfixed in 4% PFA, 2.5% glutaraldehyde in 0.1 M PB for 2 hours at room temperature and then at 4°C overnight. The sample was prepared using the BROPA (brain-wide reduced-osmium staining with pyrogallol-mediated amplification) approach and embedded by epon-812 resin (SPI-epon-812, SPI) ^46^. In specific, the sample was washed three times with 0.1 M PB for 20 minutes each time and then transferred to a 0.1 M PB solution containing 2% OsO_4_ (18451, Ted Pella) and 2.5 M formamide (47671, Sigma) for 1.5 hours at room temperature. Subsequently, the sample was treated with a 0.1 M PB solution containing 3% potassium hexacyanoferrate (60279, Sigma) for 1.5 hours and then stained with fresh 0.1 M PB solution containing 2% OsO_4_ for 1.5 hours. After washing with 0.1 M PB solution, the sample was treated with a 320 mM aqueous solution of pyrogallol (16040, Sigma) at pH 4.1 for 1.5 hours, washed with ddH_2_O and stained with a fresh 2% OsO_4_ aqueous solution for 1 hour. After washing with ddH_2_O, the sample was stained overnight at 4L using 2% uranyl acetate (02624-AB, SPI-chem) followed by heating and dying in a 50L oven for 30 minutes. After washing with ddH_2_O, the sample was dehydrated with 50%, 70%, 80%, 90%, and 100% ethanol, 1:1 ethanol: acetone, and acetone three times in sequence and each step of gradient dehydration lasted for 12 minutes. The sample was then subjected to epoxy resin penetration by treating with epoxy resin/acetone (1/1, 2/1, 3/1) and pure resin twice under negative pressure and each treatment lasted for 4-8 hours. The sample was placed in a resin-embedded plate and adjusted in position, then polymerized in a 60L oven for 24 hours. The samples were sectioned using a Leica EM UC7 ultramicrotome (Leica) with a section thickness of 70 nm and collected on silicon wafers. The images were acquired under a Focused ion beam scanning electron microscope (Crossbeam 550, Zeiss) with an accelerating voltage of 3 kV. ImageJ was used for data analysis. The mitochondrial circularity was defined by 4π_lJ_(mean area/mean perimeter^2^).

### *Romo1* knockdown in NSC34 cells

NSC34 cells which is a motor neuron-like hybrid cell line were cultured in Dulbecco’s modified Eagle’s medium (Macgene) supplemented with 10% fetal bovine serum (Gibco), 100 U/mL penicillin (Macgene), and 100 µg/mL streptomycin sulfate (Macgene) at 37°C. For *Romo1* knockdown, 150 pmol siRNA were transfected using Lipofectamine RNAiMAX (Invitrogen). The siRNAs used were a mixture of three siRNAs (*Romo1*-siRNA 1: 5’-CCAUUGGAAUGGGCAUACGAUTT-3’ (sense), 5’-AUCGUAUGCCCAUUCCAAUGGTT-3’ (antisense); *Romo1*-siRNA 2: 5’-CGUGAAGAUGGGCUUCGUCAUTT-3’ (sense), 5’-AUGACGAAGCCCAUCUUCACGTT-3’ (antisense); *Romo1*-siRNA 3: 5’-CGGCACGUUUGGCACUUUCAUTT-3’ (sense), 5’-AUGAAAGUGCCAAACGUGCCGTT-3’ (antisense)). The negative control siRNA is 5’-UUCUCCGAACGUGUCACGUTT-3’ (sense), 5’-ACGUGACACGUUCGGAGAATT-3’ (antisense). The cells were collected after 72 hours of transfection for further analysis.

### Western blot analysis

NSC34 cells were lysed with ice-cold RIPA buffer (1053+, Applygen) containing protease inhibitor cocktail (P1005, Beyotime). Supernatants were collected after centrifugation for 10 minutes at 15000 rpm and the protein concentration was measured using Rapid Gold BCA protein assay (A55861, Thermo Fisher Scientific). The protein samples were separated by 15% glycine SDS-PAGE and transferred to PVDF membranes (0.22 μm). After blocking with 5% non-fat milk in TBST (150 mM NaCl, 50 mM Tris, 0.1% Tween-20, pH 7.35) solution, membranes were incubated with primary antibodies (monoclonal anti-ROMO1 (TA505611, Thermo Fisher Scientific, 1:2000) and monoclonal anti-GAPDH (A19056, ABclonal, 1:3000)) diluted in 5% non-fat milk TBST solution overnight at 4°C. Blots were then visualized using secondary antibodies conjugated with IRDye by an Odyssey imaging system.

### Acetylcholine and choline analysis

The acetylcholine and choline contents in mouse tibialis anterior muscle or NSC34 cells were analyzed using liquid chromatography-mass spectrometry. Briefly, the tissue or NSC34 cells were rapidly isolated and washed twice in pre-cooled PBS. Then, the tissue or NSC34 cells were homogenized and sonicated in acetonitrile containing 0.1% formic acid on ice. The samples were then centrifuged at 15000 rpm for 10 minutes at 4L. The supernatant containing acetylcholine and choline was carefully aspirated and transferred to a clean EP tube for direct measurement. 2 μL pretreated sample solution was injected into the ultra-performance liquid chromatography coupled with a triple quadrupole mass spectrometry system (ACQUITY I-Class system, Waters corporation) for detection. The target compound was separated on a InfinityLab Poroshell 120 HILIC (2.7 μm, 2.1*50 mm). The mobile phases employed were acetonitrile containing 0.05% formic acid (A) and H_2_O containing 10 mM ammonium acetate (B). The gradient program was as follows: 0 minutes 10% B, 5 minutes 50% B, 6-8 minutes 60% B, 8-12 minutes 10% B. The flow rate was 0.4 ml/minutes. During the whole analysis, the column temperature was maintained at 30L and the autosampler was at 4L to avoid sample degradation. The ion source temperature and capillary voltage were kept constant and set to 150L and 2.5 kV respectively. The cone gas flow rate was 20 L/hour and the desolvation temperature was 400L. The multiple reaction monitoring parameter was listed in the following table Compound Parent (m/z) Daughter (m/z) Cone (V) Collision energy (v) mode

| Compound | Parent (m/z) | Daughter (m/z) | Cone (V) | Collision energy (v) | mode |
| --- | --- | --- | --- | --- | --- |
| L-choline | 104.1 | 45.1 | 22 | 15 | ESI+ |
| acetylcholine | 146.22 | 86.98 | 25 | 10 | ESI+ |

### Measurement of cellular ATP

Cellular ATP levels were measured using liquid chromatography-tandem mass spectrometry. The cultured NSC34 cells were gently collected and divided into two equal parts for ATP detection and protein quantitation. The collected cell samples were mixed with 200 μL 80% methanol and vortexed. The cells were then sonicated in an ice-water mixture, with a sonication time of 30 seconds followed by a 10-second pause, repeated six times. The sonicated cell suspension was centrifuged 12000 rpm for 10 minutes at 4L. The supernatant was transferred to a clean EP tube for the following measurement. 2 μL pretreated sample solution and standards were injected into Agilent 1260-Ultivo high-performance infinity liquid chromatography triple quadrupole system (Agilent, USA) for detection. ATP was separated on a InfinityLab Poroshell 120 HILIC-Z.P (2.7 μm, 2.1*100 mm). The mobile phase employed was water containing 10 mM ammonium acetate (A) and acetonitrile containing 10 mM ammonium acetate (B), both were added InfinityLab deactivator additive (Agilent part number: 5191-4506, pH=9). The gradient program was as follows: 0 minutes 88% B, 7 minutes 75% B, 7.01 minutes 40% B, 8 minutes 40% B, 8.01 minutes 88% B, 12 minutes 88% B. The flow rate was 0.6 ml/minute. The column temperature was maintained at 40°C and the autosampler was at 10°C. ATP was detected using ion transition 508>136.1 m/z on positive mode, with fragmentation fragmentor at 135V and collision energy at 10V. multiple reaction monitoring parameter data were acquired using Masshunter software 0.8.0 (Agilent, USA).

### Measurement of cellular ROS

H_2_DCFA (C6827, Invitrogen) was used to measure ROS level in NSC34 cells. The were stained with H_2_DCFA (5 μM) in Tyrode’s solution (138 mM NaCl, 3.7 mM KCl, 1.2 mM NaH_2_PO_4_, 15mM D-glucose, 20mM HEPES, and 1mM CaCl_2_, pH 7.4) for 15 minutes at 37°C, followed by washing three times with Tyrode’s solution. Images taken with Zeiss LSM 980 confocal microscopy with excitation at 488 nm and at >520 nm at room temperature. The intensity of H_2_DCFA fluorescence was analyzed using Zen Blue 3.8 and ImageJ software (US National Institutes of Health).

### Statistical analysis

Data are presented as mean ± s.e.m. of multiple biological replicates or independent experiments as indicated. Statistical analysis was performed using GraphPad Prism 8 software (GraphPad Software). When appropriate, unpaired t test, unpaired two-tailed Mann-Whitney U-Test, and two-way ANOVA with Sidak’s multiple comparison test were used. For Kaplan-Meier survival curve comparisons, Mantel-Cox log-rank analysis were used. P<0.05 was considered statistically significant.

## Acknowledgments

We thank Prof. Zhuan Zhou for providing the *ChAT-cre* mic and Dr. Li Quan for mass spectrometry analysis.

## Author contributions

Conceptualization, W.N., H.C., and X.W.; Methodology, W.N., L.L., F.X., H.H.; Investigation, W.N., Z.J., L.L., L.L., X.D.; Writing–Original Draft, W.N., L.C., and X.W.; Writing –Review & Editing, W.N., L.C., H.C., and X.W.; Funding Acquisition, W.N., H.H., H.C. and X.W.; Supervision, H.C. and X.W..

## Availability of data and materials

The dataset(s) supporting the conclusions of this article is(are) included within the article (and its additional file(s))

