## supplemental figures for "ROMO1 loss in cholinergic neurons induces mitochondrial ultrastructural damage and ALS-like neuromuscular degeneration"

**Supplementary material**

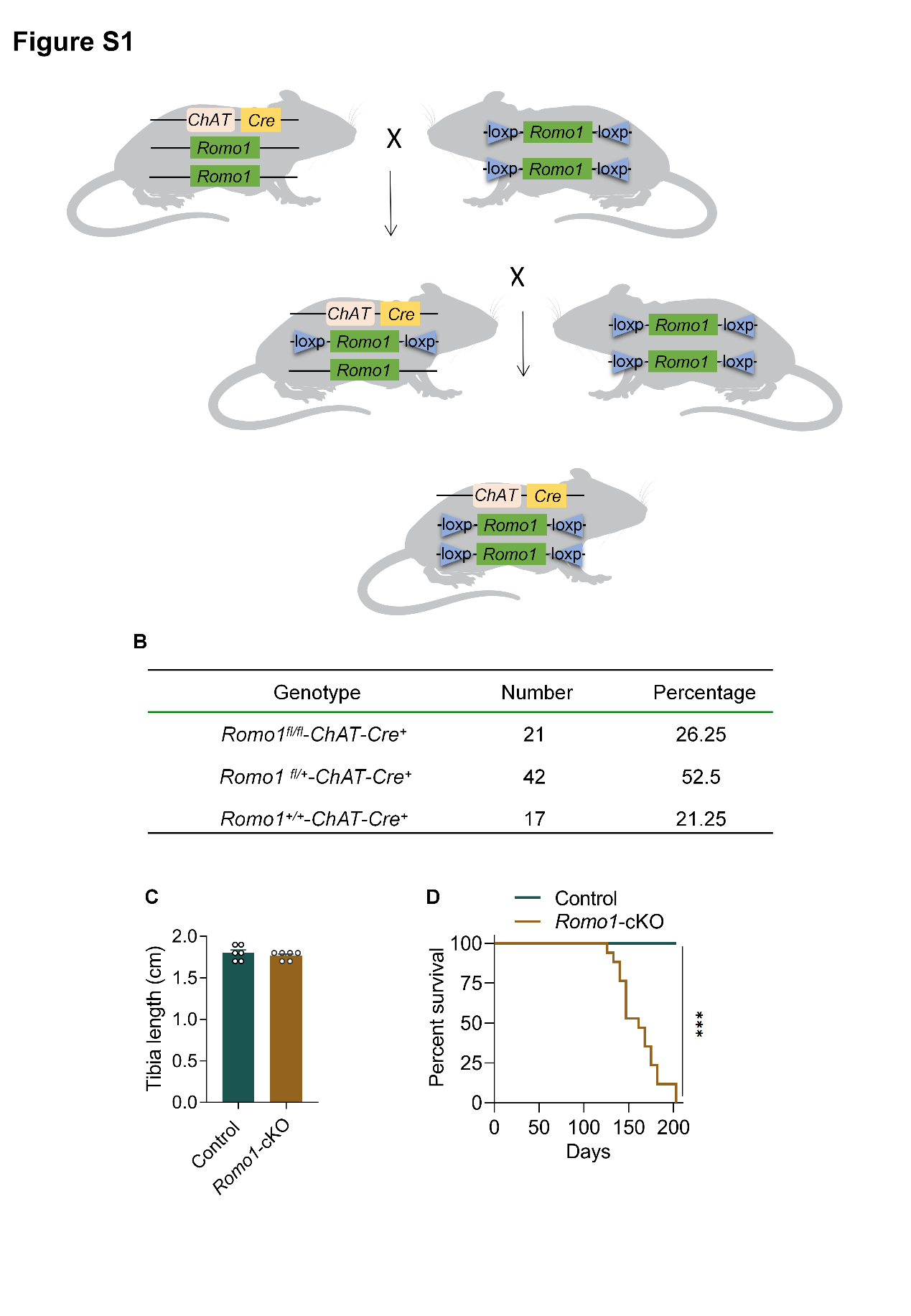

**Figure S1.** **Construction of** ***Romo1*-cKO mouse model, related to Figure 1.**

(**A**) Schematic of breeding strategy for generating *Romo1*-cKO mice. *Romo1* *flox/flox* mice were crossed with *ChAT-Cre* mice to allow cholinergic-selective deletion of *Romo1*.

(**B**) Summary of live pups recovered from *Romo1* ^+/−^ intercrossing under ChAT-Cre.

(**C**) Comparable tibia length between control and *Romo1*-cKO mice at the age of 12-month. Data were mean ± SEM. n=6 tibias for each group. Unpaired t test was used.

(**D**) Kaplan-Meier survival curves showing shorted lifespan of female *Romo1*-cKO mice after breeding. n=18 mice for each group. Mantel-Cox log-rank analysis was used. ***p < 0.001.

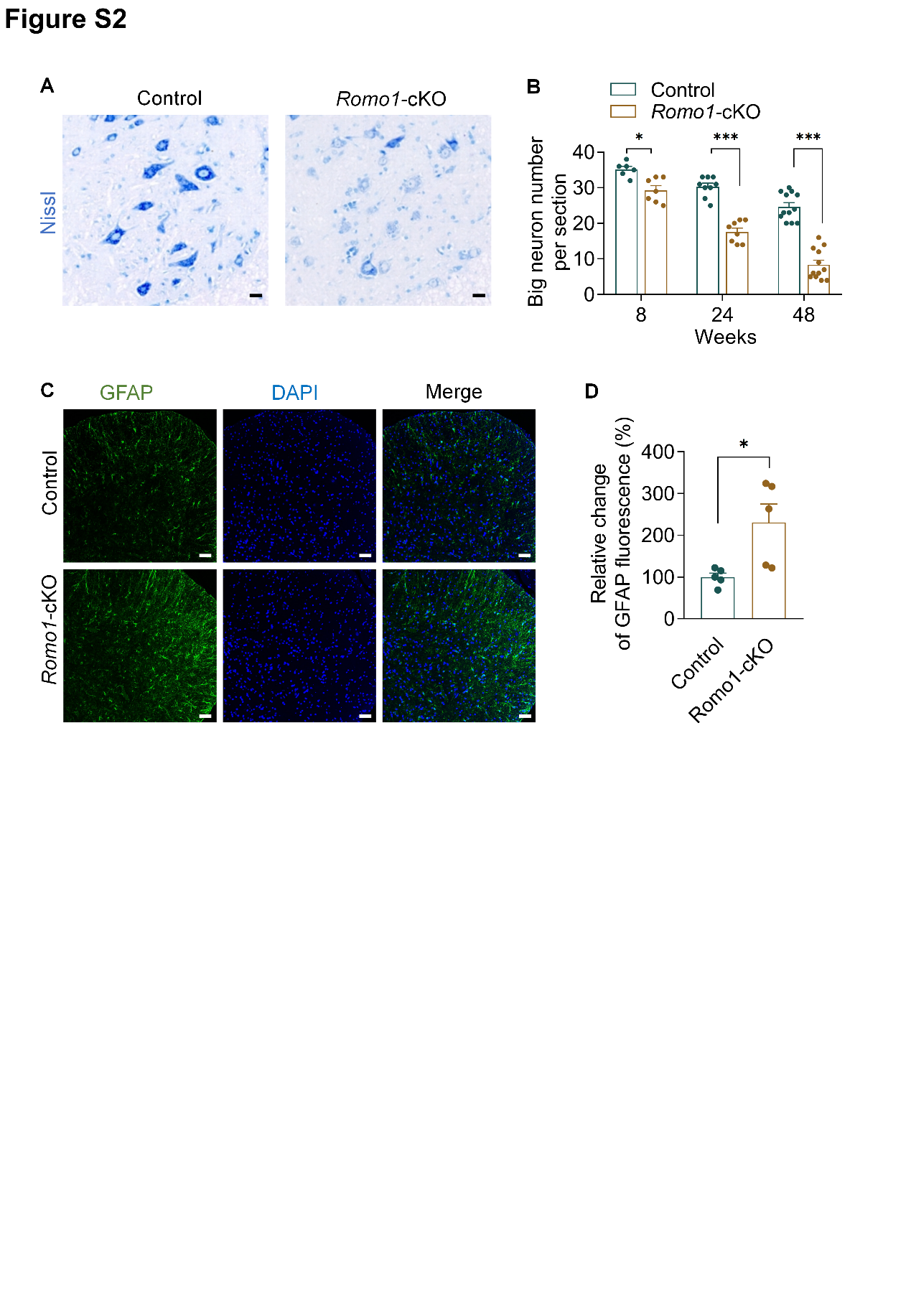

**Figure S2. Loss of big neurons and gliosis in the spinal cord of *Romo1*-cKO mice, related to Figure 2.**

(**A**) Representative Nissl staining of big neurons in the ventral horn of level 5 lumbar spinal cord isolated from *Romo1*-cKO and control mice at 48 weeks old. Scale bars: 20 μm.

(**B**) Quantification of big neuron numbers in the level 5 lumbar spinal cord at different ages of *Romo1*-cKO and control mice. Data were mean ± SEM. n=6-12 slices from 3-6 mice for each group.

(**C**) Representative images of the ventral horn in the level 5 lumbar spinal cord stained with DAPI for nuclei and GFAP for astrocytes. 8-week old *Romo1*-cKO and control mice were used. Scale bars: 50 μm.

(**D**) Quantification of GFAP-positive immunoreactivity in (C). Data were mean ± SEM. n=5 mice for each group.

For B and D, unpaired t test was used. *p < 0.05, ***p < 0.001.

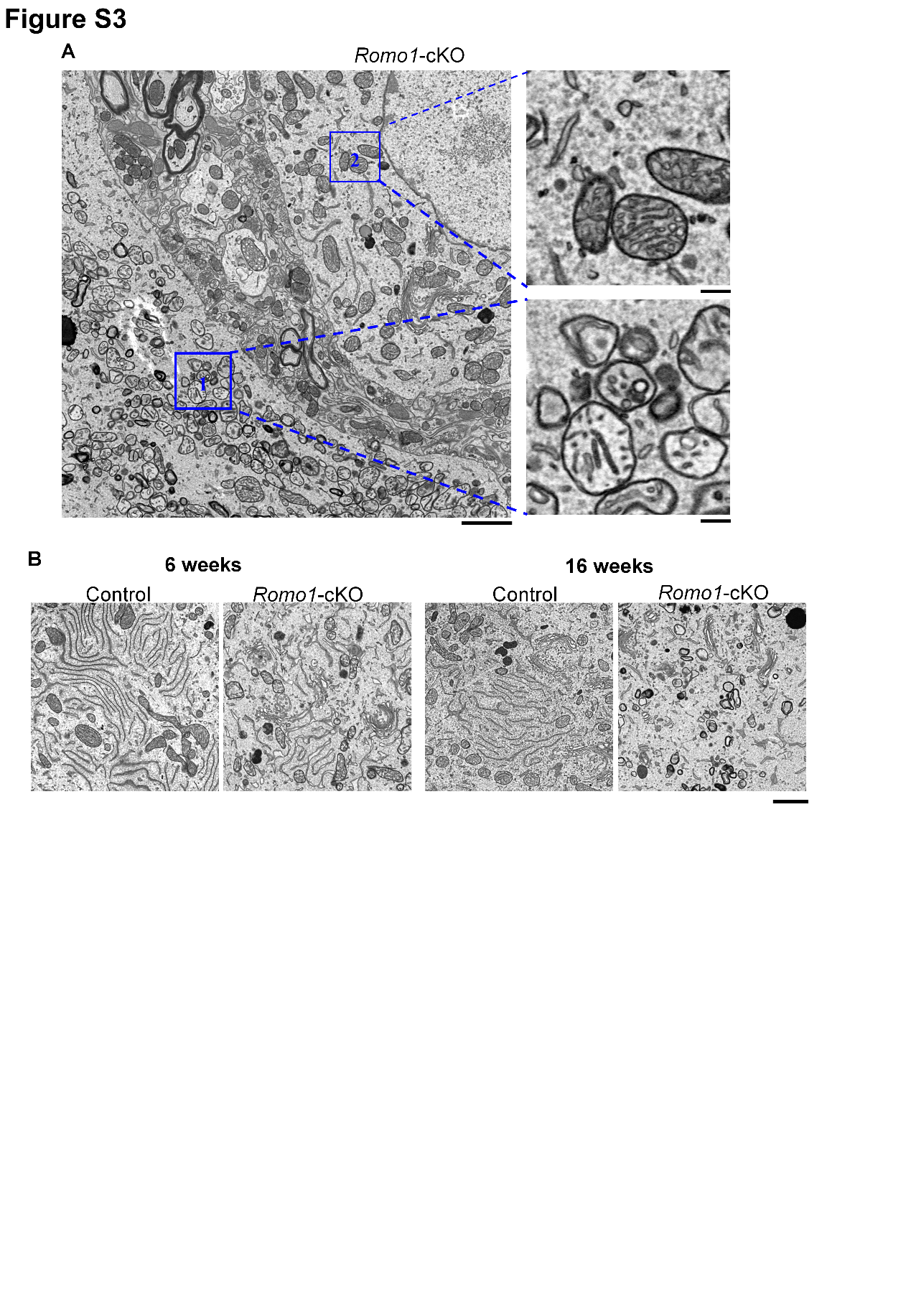

**Figure S3.** **Selective cristae deformation and ER structure disruption in *Romo1*-cKO motor neurons, related to Figure 4.**

(**A**) Representative electron micrograph of neuron somas in the level 5 lumbar spinal cord in 16-week old *Romo1*-cKO mice. Note that mitochondrial cristae were specifically disrupted in the motor neuron (1), whereas the adjacent neuron showed normal lamellar cristae structure (2). Scale bars: 2 μm (left), 300 nm (right).

(**B**) Representative electron micrographs showing disruption of ER structure in the *Romo1*-cKO motor neurons at 6- and 16-week old. Scale bar: 400 nm.

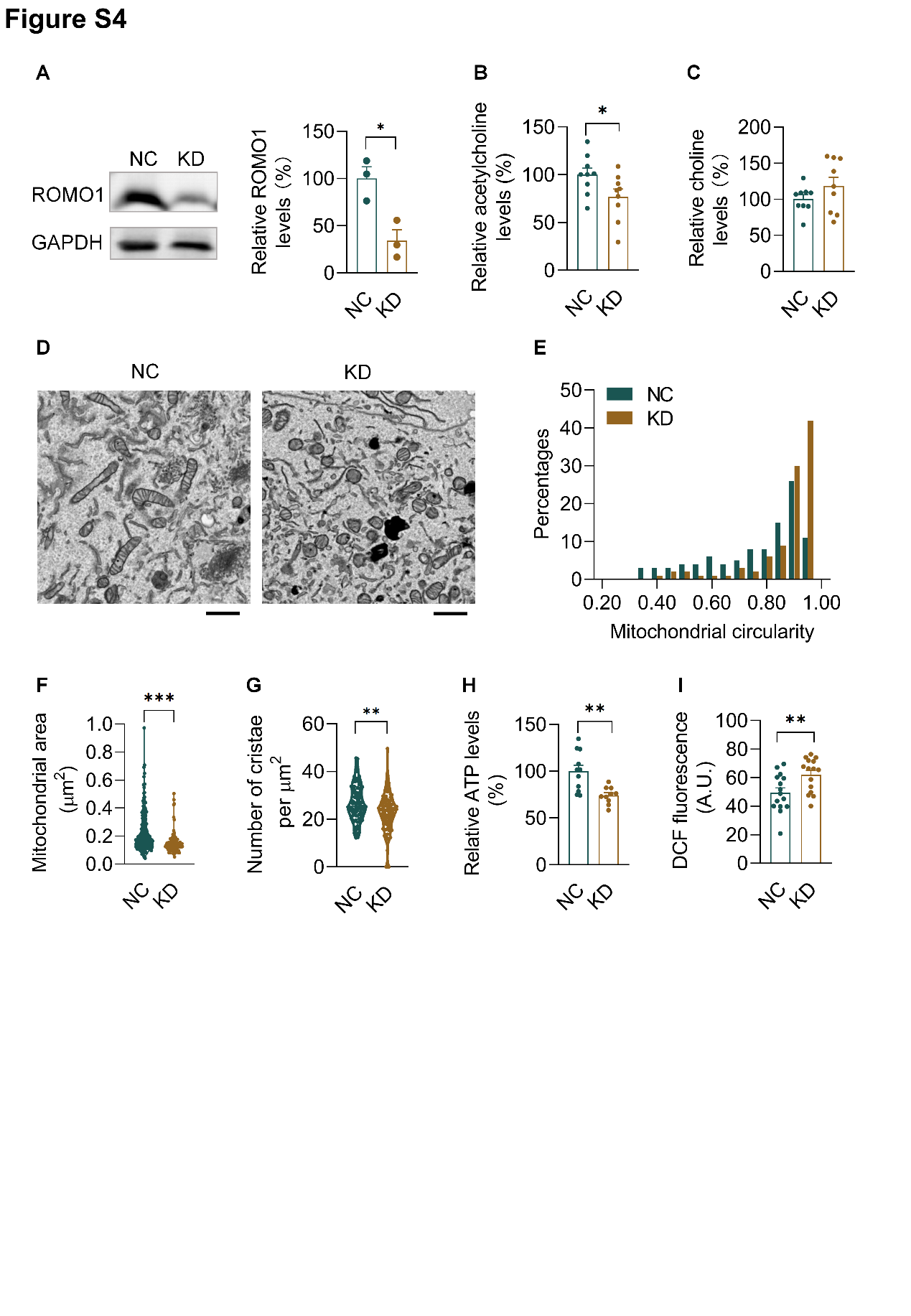

**Figure S4. ROMO1 downregulation induced structural and functional damage of mitochondria in NSC34 cells.**

(**A**) Left: Western blot showing knockdown of *Romo1* in NSC34 cells. GAPDH served as the loading control. Right: Qualification. NC, negative control. KD, *Romo1* knockdown.

(**B,C**) Effects of *Romo1* knockdown on intracellular acetylcholine (B) and choline levels (C) in NSC34 cells. n=9 dishes from 3 independent experiments for each group.

(**D**) Representative electron micrographs of NC and KD NSC34 cells. Scale bars: 1μm .

(**E**) Frequency distribution of mitochondrial circularity in NC and KD NSC34 cells. n=100 mitochondria from 3-4 cells for each group.

(**F**) Quantification of mitochondrial area in NC and KD NSC34 cells. n=100 mitochondria from 3-4 cells for each group.

(**G**) Quantification of cristae number per mitochondrion in NC and KD NSC34 cells. n=100 mitochondria from 3-4 cells for each group.

(**H**) Decreased ATP contents in KD NSC34 cells. n=10-11 dishes from 3 independent experiments for each group.

(**I**) Increased ROS levels in KD NSC34 cells. n=15 dishes from 4 independent experiments for each group.

For A-C and H-I, unpaired t test was used; for F and G, unpaired, two-tailed Mann-Whitney U-Test was used. *p < 0.05, **p < 0.01, ***p < 0.001.

**Movie S1. Locomotor activities of 20-week old control and *Romo1*-cKO mice. Note that *Romo1*-cKO mouse displays hindlimb paralysis at this age.**
